# SigProfilerMatrixGenerator: a tool for visualizing and exploring patterns of small mutational events

**DOI:** 10.1101/653097

**Authors:** Erik N. Bergstrom, Mi Ni Huang, Uma Mahto, Mark Barnes, Michael R. Stratton, Steven G. Rozen, Ludmil B. Alexandrov

## Abstract

**Background:** Cancer genomes are peppered with somatic mutations imprinted by different mutational processes. The mutational pattern of a cancer genome can be used to identify and understand the etiology of the underlying mutational processes. A plethora of prior research has focused on examining mutational signatures and mutational patterns from single base substitutions and their immediate sequencing context. We recently demonstrated that further classification of small mutational events (including substitutions, insertions, deletions, and doublet substitutions) can be used to provide a deeper understanding of the mutational processes that have molded a cancer genome. However, there has been no standard tool that allows fast, accurate, and comprehensive classification for all types of small mutational events

**Results:** Here, we present SigProfilerMatrixGenerator, a computational tool designed for optimized exploration and visualization of mutational patterns for all types of small mutational events. SigProfilerMatrixGenerator is written in Python with an R wrapper package provided for users that prefer working in an R environment. SigProfilerMatrixGenerator produces fourteen distinct matrices by considering transcriptional strand bias of individual events and by incorporating distinct classifications for single base substitutions, doublet base substitutions, and small insertions and deletions. While the tool provides a comprehensive classification of mutations, SigProfilerMatrixGenerator is also faster and more memory efficient than existing tools that generate only a single matrix.

**Conclusions:** SigProfilerMatrixGenerator provides a standardized method for classifying small mutational events that is both efficient and scalable to large datasets. In addition to extending the classification of single base substitutions, the tool is the first to provide support for classifying doublet base substitutions and small insertions and deletions. SigProfilerMatrixGenerator is freely available at https://github.com/AlexandrovLab/SigProfilerMatrixGenerator with an extensive documentation at https://osf.io/s93d5/wiki/home/.

## Background

Analysis of somatic mutational patterns is a powerful tool for understanding the etiology of human cancers [1]. The examination of mutational patterns can trace its origin to seminal studies that evaluated the patterns of mutations imprinted in the coding regions of *TP53* [2], the most commonly mutated gene in human cancer [3]. These early reports were able to identify characteristic patterns of single point substitutions imprinted due to smoking tobacco cigarettes, exposure to ultraviolet light, consumption of aflatoxin, intake of products containing aristolochic acid, amongst others [4-7]. The advent of massively parallel sequencing technologies [8] allowed cheap and efficient evaluation of the somatic mutations in a cancer genome. This provided an unprecedented opportunity to examine somatic mutational patterns by sequencing multiple cancer-associated genes, by sequencing all coding regions of the human genome (i.e., usually referred to as whole-exome sequencing), or even by interrogating the complete sequence of a cancer genome (i.e., an approach known as whole-genome sequencing).

Examinations of mutational patterns from whole-genome and whole-exome sequenced cancers confirmed prior results derived from evaluating the mutations in the coding regions of *TP53* [9]. For example, the cancer genome of a lung cancer patient with a long history of tobacco smoking was peppered with somatic mutations exhibiting predominately cytosine to adenine single base substitutions [10]; the same mutational pattern was previously reported by examining mutations in *TP53* in lung cancers of tobacco smokers [4, 11]. In addition to confirming prior observations, whole-exome and whole-genome sequencing data provided a unique opportunity for identifying all of the mutational processes that have been active in the lineage of a cancer cell [12]. By utilizing mathematical modelling and computational analysis, we previously created the concept of mutational signatures and provided tools for deciphering mutational signatures from massively parallel sequencing data [13]. It should be noted that a mutational signature is mathematically and conceptually distinct from a mutational pattern of a cancer genome. While a mutational pattern of a cancer genome can be directly observed from sequencing data, a mutational signature is, in most cases, not directly observable. Rather, a mutational signature corresponds to a mathematical abstraction (i.e., a probability mass function) derived through a series of numerical approximations. From a biological perspective, a mutational signature describes a characteristic set of mutation types reflecting the activity of endogenous and/or exogenous mutational processes [12]. By examining the directly observed mutational patterns of thousands of cancer genomes, we were able to identify 49 single point substitution, 11 doublet base substitution, and 17 small insertion and deletion signatures [14] in human cancer and to propose a putative etiology for a number of these signatures.

Since we presented the very first bioinformatics framework for deciphering mutational signatures in cancer genomes [13, 15], a number of computational tools have been developed for the analysis of mutational signatures (recently reviewed in [16]). All of these tools perform a matrix factorization or leverage an approach mathematically equivalent to a matrix factorization. As such, each of these tools directly or indirectly requires generating a correct initial input matrix for subsequent analysis of mutational signatures. In principle, creating an input matrix can be examined as a transformation of the mutational catalogues of a set of cancer genomes to a matrix where each sample has a fixed number of mutation classes (also, known as mutation channels). The majority of existing tools have focused on analyzing data using 96 mutation classes corresponding to a single base substitution and the 5’ and 3’ bases immediately adjacent to the mutated substitution. While this simple classification has proven powerful, additional classifications are required to yield greater understanding of the operative mutational processes in a set of cancer genomes [12].

Here, we present SigProfilerMatrixGenerator, a computational package that allows efficient exploration and visualization of mutational patterns. SigProfilerMatrixGenerator is written in Python with an R wrapper package provided for users that prefer working in an R environment. The tool can read somatic mutational data in most commonly used data formats (i.e., VCF, MAF, etc.) and it provides support for analyzing all types of small mutational events: single bases substitutions, doublet base substitutions, and small insertions and deletions. SigProfilerMatrixGenerator generates fourteen distinct matrices including ones with extended sequencing context and transcriptional strand bias, while providing publication ready visualization for the majority of these matrices. Further, the tool is the first to provide standard support for the classification of small insertions and deletions as well as the classification of doublet base substitutions that were recently used to derive the next generation of mutational signatures [14]. While SigProfilerMatrixGenerator provides much more functionality (Table 1), in almost all cases, it is more computationally efficient than existing approaches. Lastly, SigProfilerMatrixGenerator comes with extensive Wiki-page documentation and can be easily integrated with existing packages for analysis of mutational signatures.

**Table 1:** Matrix generation and visualization functionality of six commonly used tools. **M** corresponds to providing functionality to only generate a mutational matrix; **MP** corresponds to providing functionality to both generate and plot a mutational matrix. **\*** indicates that a tool can perform only one of the actions in a single run; for example, Helmsman can either generate a 96 or a 1536 mutational matrix but not both in a single run.

| Tool | SBS |  |  |  |  |  | ID |  |  |  | DBS |  |  |  |
| --- | --- | --- | --- | --- | --- | --- | --- | --- | --- | --- | --- | --- | --- | --- |
|  | 6 | 24 | 96 | 384 | 1536 | 6144 | 28 | 83 | 415 | 8628 | 78 | 186 | 1248 | 2976 |
| SigProfilerMatrixGenerator<br>Language: Python & R | MP | MP | MP | MP | MP | M | MP | MP | MP | M | MP | MP | M | M |
| Helmsman [31]<br>Language: Python |  |  | M* |  | M* |  |  |  |  |  |  |  |  |  |
| deconstructSigs [27]<br>Language: R |  |  | MP |  |  |  |  |  |  |  |  |  |  |  |
| mafTools [28]<br>Language: R | MP |  | MP |  |  |  |  |  |  |  |  |  |  |  |
| SomaticSignatures [29]<br>Language: R |  |  | MP* |  | M* |  |  |  |  |  |  |  |  |  |
| signeR [30]<br>Language: R |  |  | MP* |  | M* |  |  |  |  |  |  |  |  |  |

## Implementation

### Classification of Single Base Substitutions (SBSs)

A single base substitution (SBS) is a mutation in which a single DNA base-pair is substituted with another single DNA base-pair. An example of an SBS is a **C:G** base-pair mutating to an **A:T** base-pair; this is usually denoted as a **C:G>A:T**. The most basic classification catalogues SBSs into six distinct categories, including: C:G>A:T, C:G>G:C, C:G>T:A, T:A>A:T, T:A>C:G, and T:A>G:C. In practice, this notation has proven to be bulky and, in most cases, SBSs are referred to by either the purine or the pyrimidine base of the Watson-Crick base-pair. Thus, one can denote a **C:G>A:T** substitution as either a **C>A** mutation using the pyrimidine base or as a **G>T** mutation using the purine base. While all three notations are equivalent, prior research on mutational signatures [13, 15, 17] has made the pyrimidine base of the Watson-Crick base-pair a community standard. As such, the most commonly used SBS-6 classification of single base substitutions can be written as: C>A, C>G, C>T, T>A, T>C, and T>G. The classification SBS-6 should not be confused with signature SBS6, a mutational signature attributed to microsatellite instability [15].

The simplicity of the SBS-6 classification allows capturing the predominant mutational patterns when only a few somatic mutations are available. As such, this classification was commonly used in analyzing mutational patterns derived from sequencing *TP53* [4, 11]. The SBS-6 classification can be further expanded by taking into account the base-pairs immediately adjacent 5’ and 3’ to the somatic mutation. A commonly used classification for analysis of mutational signatures is SBS-96, where each of the classes in SBS-6 is further elaborated using one base adjacent at the 5’ of the mutation and one base adjacent at the 3’ of the mutation. Thus, for a C>A mutation, there are sixteen possible trinucleotide (4 types of 5’ base ∗ 4 types of 3’ base): A<u>C</u>A>A<u>A</u>A, A<u>C</u>C>A<u>A</u>C, A<u>C</u>G>A<u>A</u>G, A<u>C</u>T>A<u>A</u>T, C<u>C</u>A>C<u>A</u>A, C<u>C</u>C>C<u>A</u>C, C<u>C</u>G>C<u>A</u>G, C<u>C</u>T>C<u>A</u>T, G<u>C</u>A>G<u>A</u>A, G<u>C</u>C>G<u>A</u>C, G<u>C</u>G>G<u>A</u>G, G<u>C</u>T>G<u>A</u>T, T<u>C</u>A>T<u>A</u>A, T<u>C</u>C>T<u>A</u>C, T<u>C</u>G>T<u>A</u>G, and T<u>C</u>T>T<u>A</u>T (mutated based is underlined). Each of the six single base substitutions in SBS-6 has sixteen possible trinucleotides resulting in a classification with 96 possible channels (Fig 1A). In this notation, the mutated base is underlined and the pyrimidine base of the Watson-Crick base-pair is used to refer to each SBS. Please note that using the purine base of the Watson-Crick base-pair for classifying mutation types will require taking the reverse complement sequence of each of the classes of SBS-96. For example, A<u>C</u>G:T<u>G</u>C>A<u>A</u>G:T<u>T</u>C can be written as A<u>C</u>G>A<u>A</u>G using the pyrimidine base and as C<u>G</u>T>C<u>T</u>T using the purine base (i.e., the reverse complement sequence of the pyrimidine classification). Similarly, an A<u>G</u>C:T<u>C</u>G>A<u>A</u>C:T<u>T</u>G mutation can be written as A<u>G</u>C>A<u>A</u>C using the purine base and G<u>C</u>T>G<u>T</u>T using the pyrimidine base (i.e., the reverse complement sequence of the purine classification). In principle, somatic mutations are generally reported based on the reference strand of the human genome thus requiring converting to either the purine or the pyrimidine base of the Watson-Crick base-pair. Prior work on mutational signatures [13–15, 17] has established the pyrimidine base as a standard for analysis of somatic mutational patterns.

**Figure 1:**
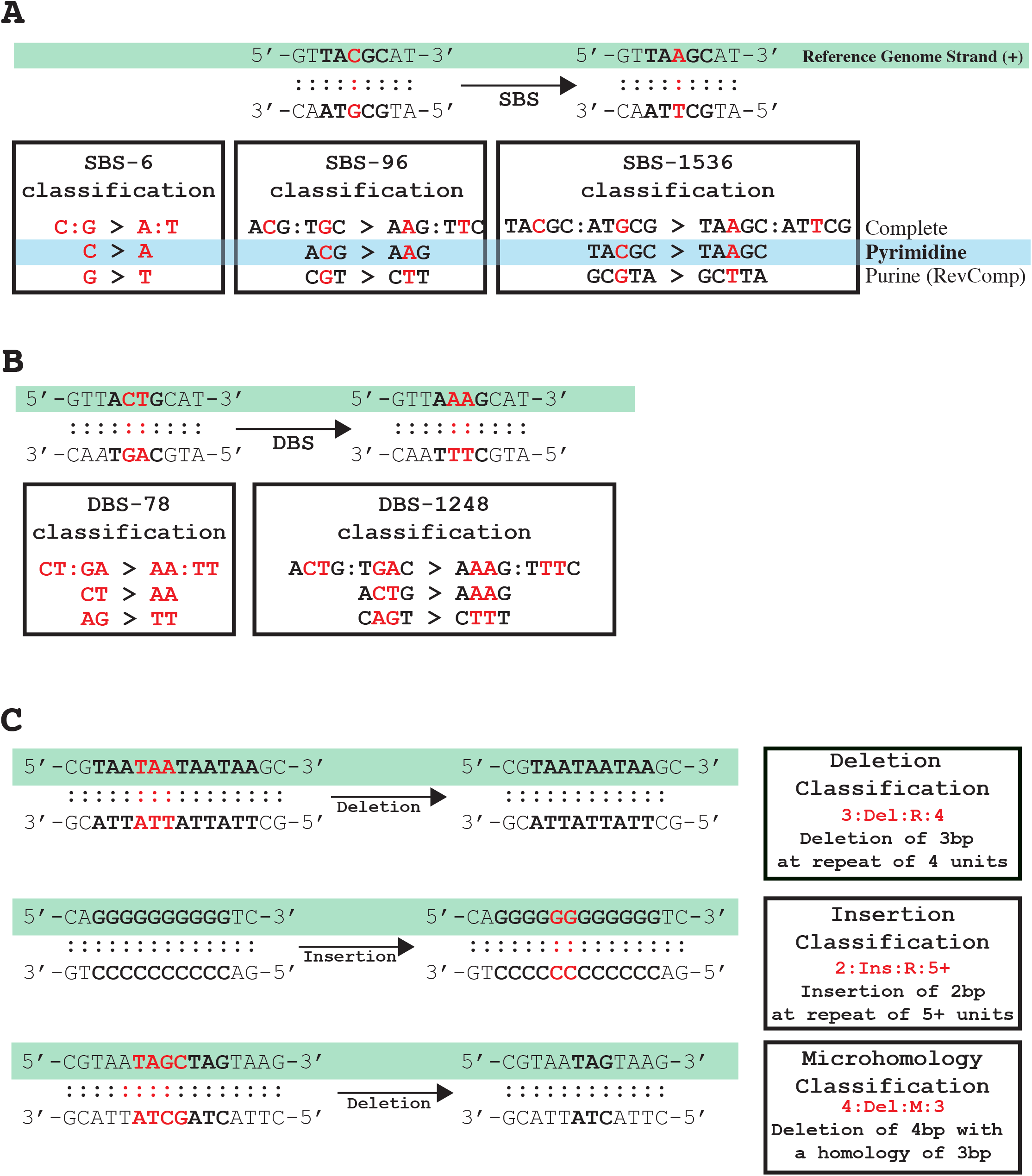
Classifications of single base substitutions, doublet base substitutions, and indels. ***A)*** Classification of single base substitutions (SBSs). The complete classification of an SBS includes both bases in the Watson-Crick base-pairing. To simplify this notation, one can use either the purine or the pyrimidine base. SigProfilerMatrixGenerator uses as a standard the pyrimidine classification. ***B)*** Classification of doublet base substitutions (DBSs). The complete classification of a DBS includes bases on both strands. To simplify this notation, in most cases, SigProfilerMatrixGenerator uses the maximum number of pyrimidines. ***C)*** Classification of small insertions and deletions. The complete classification includes the length of the indel and the number of repeated units surrounding the sequence. For deletions at microhomologies, the length of the homology, rather than the number of repeat units surrounding the indel, is used in the classification.

The SBS-96 has proven particularly useful for analysis of data from both whole-exome and whole-genome sequencing data [17]. This classification is both simple enough to allow visual inspection of mutational patterns and yet sufficiently complicated for separating different sources of the same type of an SBS. For example, mutational signatures analysis has identified at least 15 distinct patterns of C>T mutations each of which has been associated with different mutational processes (e.g., exposure to ultraviolet light [18], activity of the APOBEC family of deaminases [19], failure of base excision repair [20], etc.). SBS-96 can be further elaborated by including additional sequencing context. Simply by including additional 5’ and 3’ adjacent context, one can increase the resolution. For example, considering two bases 5’ and two bases 3’ of a mutation results in 256 possible classes for each SBS (16 types of two 5’ bases ∗ 16 types of two 3’ bases). Each of the six single base substitutions in SBS-6 has 256 possible pentanucleotides resulting in a classification with 1536 possible channels. Since we first introduced SBS-1536 [13], this classification has found limited use in analysis of mutational patterns. The increased number of mutational channels requires a large number of somatic mutations, which can be generally found only in whole-genome sequenced cancer exhibiting a high mutational burden (usually >2 mutations per MB). Nevertheless, SBS-1536 has been used to further elaborate the mutational patterns exhibited by several mutagenic processes, for example, the aberrant activity of DNA polymerase epsilon [14] or the ectopic action of the APOBEC family of cytidine deaminases [13, 14].

SigProfilerMatrixGenerator provides matrix generation support for SBS-6, SBS-96, and SBS-1536 using the commonly accepted pyrimidine base of the Watson-Crick base-pair. Further, the tool allows interrogation of transcriptional strand bias for each of these classifications and provides a harmonized visualization for all three matrices.

### Classification of Doublet Base Substitutions (DBSs)

A doublet base substitution (DBS) is a somatic mutation in which a set of two adjacent DNA base-pairs is simultaneously substituted with another set of two adjacent DNA base-pairs. An example of a DBS is a set of **CT:GA** base-pairs mutating to a set of **AA:TT** base-pairs, which is usually denoted as **CT:GA>AA:TT** (Fig 1B). It should be noted that a **CT:GA>AA:TT** mutation can be equivalently written as either a CT>AA mutation or an AG>TT mutation (note that AG>TT is the reverse complement of CT>AA). Similar to the SBSs, the complete notation for DBS has proven bulky. As such, we have previously defined a canonical set of DBSs and used this set to interrogate both mutational patterns and mutational signatures [14]. In this canonical set, DBSs are referred to using the maximum number of pyrimidine nucleotides of the Watson-Crick base-pairs; for example, an AA:TT>GT:CA mutation is usually denoted as TT>AC as this notation contains three pyrimidine nucleotides rather than the alternative AA>GT notation, which contains only a single pyrimidine nucleotide. There are several DBSs with the equivalent number of pyrimidine nucleotide in each context (e.g., AA:TT>CC:GG), in such cases, one of these notations was selected. Further, it should be noted, that some DBSs are palindromic. For example, an AT:TA>CG:GC can be written only as AT>CG since the reverse complement of 5’-AT-3’>5’-CG-3’ is again 5’-AT-3’>5’-CG-3’. Overall, the basic classification catalogues DBSs into 78 distinct categories denoted as the DBS-78 matrix (Supplementary Table 1).

While the prevalence of DBSs in a cancer genome is relatively low, on average a hundred times less than SBSs [14], we have previously demonstrated that a doublet base substitution is not two single base substitutions occurring simply by chance next to one another [14]. While such events are possible, across most human cancers, they will account for less than 0.1% of all observed DBSs [14]. Further, certain mutational processes have been shown to specifically generate high levels of DBSs. A flagship example is the exposure to ultraviolet light, which causes large numbers of CC>TT mutations in cancers of the skin [5]. Other notable examples are DBSs accumulating due to defects in DNA mismatch repair [14], exposure to platinum chemotherapeutics [21], tobacco smoking [22], and many others [14].

Similar to the classification of SBSs, we can expand the characterization of DBS mutations by considering the 5’ and 3’ adjacent contexts. By taking one base on the 5’ end and one base on the 3’ end of the dinucleotide mutation, we establish the DBS-1248 context. For example, a CC>TT mutation has 16 possible tetranucleotides: A<u>CC</u>A>A<u>TT</u>A, A<u>CC</u>C>A<u>TT</u>C, A<u>CC</u>G>A<u>TT</u>G, A<u>CC</u>T>A<u>TT</u>T, C<u>CC</u>A>C<u>TT</u>A, C<u>CC</u>C>C<u>TT</u>C, C<u>CC</u>G>C<u>TT</u>G, C<u>CC</u>T>C<u>TT</u>T, G<u>CC</u>A>G<u>TT</u>A, G<u>CC</u>C>G<u>TT</u>C, G<u>CC</u>G>G<u>TT</u>G, G<u>CC</u>T>G<u>TT</u>T, T<u>CC</u>A>T<u>TT</u>A, T<u>CC</u>C>T<u>TT</u>C, T<u>CC</u>G>T<u>TT</u>G, and T<u>CC</u>T>T<u>TT</u>T (mutated bases are underlined). With seventy-eight possible DBS mutations having sixteen possible tetranucleotides each, this context expansion results in 1,248 possible channels denoted as the DBS-1248 context. While this classification is provided as part of SigProfilerMatrixGenerator, it has yet to be thoroughly leveraged for analysis of mutational patterns. Further, it should be noted that for most samples, the low numbers of DBSs in a single sample will make the DBS-1248 classification impractical. Nevertheless, we expect that this classification will be useful for examining hypermutated and ultra-hypermutated human cancers.

SigProfilerMatrixGenerator generates matrices for DBS-78 and DBS-1248 by predominately using the maximum pyrimidine context of the Watson-Crick base-pairs. The matrix generator also supports the incorporation of transcriptional strand bias with an integrated display of the DBS-78 mutational patterns.

### Classification of Small Insertions and Deletions (IDs)

A somatic insertion is an event that has incorporated an additional set of base-pairs that lengthens a chromosome at a given location. In contrast, a somatic deletion is an event that has removed a set of existing base-pairs from a given location of a chromosome. Collectively, when these insertions and deletions are short (usually <100 base-pairs), they are commonly referred as small insertions and deletions (often abbreviated as indels). In some cases, indels can be complicated events in which the observed result is both a set of deleted base-pairs and a set of inserted base-pairs. For example, 5’-ATCCG-3’ mutating to 5’-ATAAAG-3’ is a deletion of CC:GG and an insertion of AAA:TTT. Such events are usually annotated as complex indels.

Indel classification is not a straightforward task and it cannot be performed analogously to SBS or DBS classifications, where the immediate sequencing context flanking each mutation was utilized to subclassify these mutational events. For example, determining the flanking sequences for deleting (or inserting) a cytosine from the sequence 5’-ATCCCCCCG-3’ is not possible as one cannot unambiguously identify which cytosine has been deleted. We recently developed a novel way to classify indels and used this classification to perform the first pan-cancer analysis of indel mutational signatures (Supplementary Table 2) [14]. More specifically, indels (IDs) were classified as single base-pair events or longer events. A single base-pair event can be further subclassified as either a **C:G** or a **T:A** indel; usually abbreviated based on the pyrimidine base as a **C** or a **T** indel. The longer indels can also be subclassified based on their lengths: 2bp, 3bp, 4bp, and 5+bp. For example, if the sequence ACA is deleted from 5’-ATTACA[ACA]GGCGC-3’ we denote this as a deletion with length 3. Similarly, if a genomic region mutates from 5’-ATTACAGGCGC-3’ to 5’-ATTACA**CCTG**GGCGC-3’, this will be denoted as an insertion with length 4 (Fig 1C).

Indels were further subclassified into ones at repetitive regions and ones with microhomologies (i.e., partial overlap of an indel). Note that microhomologies are not defined for indels with lengths of 1bp as partial overlaps are not possible. For indels with lengths of 1bp, the subclassification relied on repetitive regions that are stretches of the same base-pair referred to as homopolymers. The repeat sizes of insertions were subclassified based on their sizes of 0bp, 1bp, 2bp, 3bp, 4bp, 5+bp; while the repeat sizes of deletions were subclassified as 1bp, 2bp, 3bp, 4bp, 5bp, 6+bp (note that one cannot have a deletion with a repeat size of 0bp). For example, if the sequence ACA is deleted from 5’-ATTACA[ACA]GGCGC-3’, this will be denotated as a deletion with length 3 at a repeat unit of 2 since there are two adjacent copies of ACAACA and only one of these copies has been deleted. Similarly, if a genomic region mutates from 5’-ATTACAGGCGC-3’ to 5’-ATTACA**CCTG**GGCGC-3’, this will be denoted as an insertion with length 4 at a repeat unit of 0 since the adjacent sequences are not repeated.

In addition to classifying indels as ones occurring at repetitive regions, a classification was performed to identify the long indels with microhomologies (i.e., partially overlapping sequences). Since almost no insertions with microhomologies were identified across more than 20,000 human cancers [14], this classification was limited to long deletions at microhomologies. Microhomologies were classified based on the length of the short identical sequence of bases adjacent to the variation. For example, if TAGTC is deleted from the sequence 5’-ACCCA[TAGTC]TAGTAGCGGC-3’, this will be classified as a deletion of length five occurring at a microhomology site of length four because of the identical sequence TAGT located at the 3’ end of the deletion. Similarly, if TAGTC is deleted from the sequence 5’-ACCCAGTC[TAGTC]AAGCGGC-3’, this will also be classified as a deletion of length five occurring at a microhomology site of length four because of the identical sequence AGTC located at the 5’ end of the deletion. The classification does not distinguish (i.e., subclassify) between 3’ and 5’ microhomologies since these tend to be dependent on the mutation calling algorithms. For example, 5’-ACCCA[TAGTC]TAGTAGCGGC-3’ is the same event as 5’-ACCCATAG[TCTAG]CGGC-3’ since in both cases a 5bp sequence is deleted from a reference sequence 5’-ACCCATAGTCTAGTAGCGGC-3’and the result is 5’-ACCCATAGCGGC-3’. While somatic mutation callers may report different indels, our classification will annotate these indels as exactly the same mutational event.

The classification of small insertions and deletions was developed to reflect previously observed indel mutational processes. More specifically, the large numbers of small insertions and deletions at repetitive regions were observed in micro-satellite unstable tumors [23] as well as the large numbers of deletions were observed in tumors with deficient DNA double-strand break repair by homologous recombination [24]. Our classification was previously used to identify 17 indel signatures across the spectrum of human cancers [14]. SigProfilerMatrixGenerator allows generation of multiple mutational matrices of indels including ID-28 and ID-83. Importantly, the tool also generates an ID-8628 matrix that extends the ID-83 classification by providing complete information about the indel sequence for indels at repetitive regions with length of less than 6bp. While SigProfilerMatrixGenerator provide this extensive indel classification, ID-8628 has yet to be thoroughly utilized for analysis of indel mutational patterns. Further, it should be noted that for most samples, the low numbers of indels in a single sample will make the ID-8628 classification impractical. Nevertheless, we expect that this classification will be useful for examining cancers with large numbers of indels and especially ones with deficient DNA repair. The matrix generator also supports the incorporation of transcriptional strand bias for ID-83 and the generation of plots for most of the indel matrices.

### Incorporation of Transcription Strand Bias (TSB)

The mutational classifications described above provide a detailed characterization of mutational patterns of single base substitutions, doublet base substitutions, and small insertions and deletions. Nevertheless, these classifications can be further elaborated by incorporating additional features. Strand bias is one commonly used feature that we and others have incorporated in prior analyses [13, 15, 17]. While one cannot distinguish the strand of a mutation, one expects that mutations from the same type will be equally distributed across the two DNA strands. For example, given a mutational process that causes purely C:G>T:A mutations and a long repetitive sequence 5’-CGCGCGCGCGCGCGCGCCG-3’ on the reference genome, one would expect to see an equal number of C>T and G>A mutations. However, in many cases an asymmetric number of mutations are observed due to either one of the strands being preferentially repaired or one of the strands having a higher propensity for being damaged. Common examples of strand bias are transcription strand bias in which transcription-couple nucleotide excision repair (TC-NER) fixes DNA damage on one strand as part of the transcriptional process [25] and replicational strand bias in which the DNA replication process may result in preferential mutagenesis of one of the strands [26]. Strand bias can be measured by orienting mutations based on the reference strand. In the above-mentioned example, observing exclusively C>A mutations (and no G>A mutations) in the reference genome sequence 5’-CGCGCGCGCGCGCGCGCCG-3’ may mean that: *(i)* the guanine on the reference strand is protected; *(ii)* the cytosine on the reference strand is preferentially damaged; *(iii)* the guanine on the non-reference strand is preferentially damaged; *(iv)* the cytosine on the non-reference strand is protected; or *(v)* a combination of the previous four examples. In principle, a strand bias reveals additional strand-specific molecular mechanisms related to DNA damage, repair, and mutagenesis.

SigProfilerMatrixGenerator provides a standard support for examining transcriptional strand bias for single base substitutions, doublet base substitutions, and small indels. The tool evaluates whether a mutation occurs in the transcribed or the non-transcribed regions of well-annotated protein coding genes of a reference genome. Mutations found in the transcribed regions of the genome are further subclassified as: *(i)* transcribed, *(ii)* un-transcribed, *(iii)* bi-directional, or *(iv)* unknown. In all cases, mutations are oriented based on the reference strand and their pyrimidine context.

To sub-classify mutations based on their transcriptional strand bias, we consider the pyrimidine orientation with respect to the locations of well-annotated protein coding genes on a genome. For instance, when the coding strand (i.e., the strand containing the coding sequence of a gene; also known as the un-transcribed strand) matches the reference strand, a T:A>A:T will be reported as an untranscribed T>A (abbreviated as **U:T>A**; Fig. 2). In this case, the template strand (i.e., the strand NOT containing the coding sequence of a gene; also known as the transcribed strand) will be complementary to the reference strand and a G:C>C:G mutation will be reported as a transcribed C>G (abbreviated as **T:C>G**; Fig. 2). In rare cases, both strands of a genomic region code for a gene. Such mutations are annotated as bidirectional based on their pyrimidine context. For example, both a T:A>C:G and a A:T>G:C mutations in regions of bidirectional transcription will both be annotated as a bidirectional T>C (abbreviated as **B:T>C**). The outlined notations are applicable when describing mutations that are located within the transcribed regions of the genome. When a mutation is located outside of these regions, it will be classified as non-transcribed. For example, both a C:G>T:A and a G:C>A:T mutations in non-transcribed regions will be annotated as a non-transcribed C>T (abbreviated as **N:C>T**).

**Figure 2:**
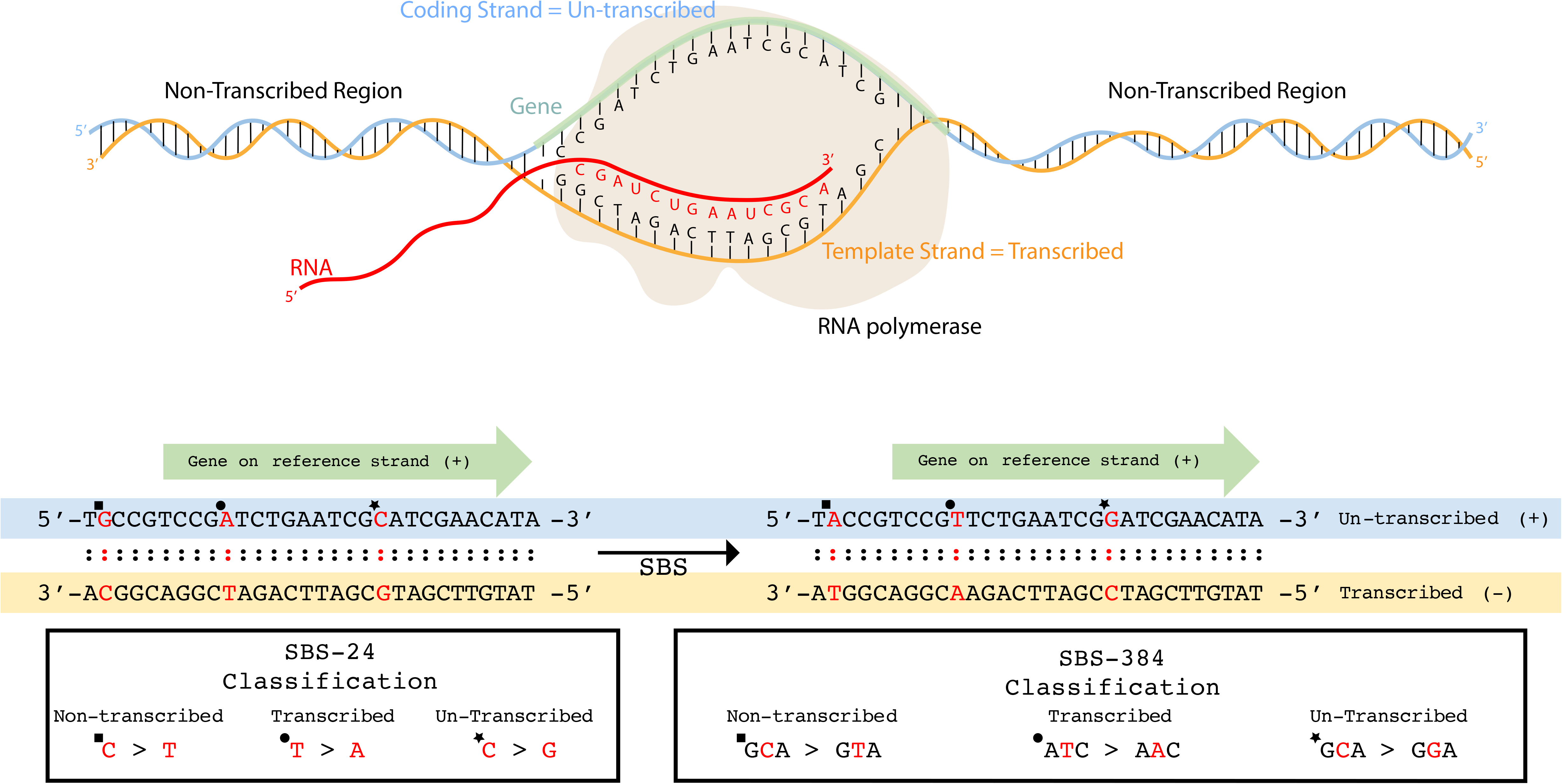
Classifications of transcriptional strand bias. ***A)*** RNA polymerase uses the template strand to transcribe DNA into RNA. The strand upon which the gene is located is referred to as the coding strand. All regions outside of the footprint of a gene are referred to as non-transcribed regions. ***B)*** Single point substitutions are oriented based on their pyrimidine base and the strand of the reference genome. When a gene is found on the reference strand an A:T>T:A substitution in the footprint of the gene is classified as transcribed T>A (example indicated by circle) while a C:G>G:C substitution in the footprint of the gene is classified as un-transcribed C>G (example indicated by star). Mutations outside of the footprints of genes are classified as non-transcribed (example indicated by square). Classification of single base substitutions is shown both in regard to SBS-24 and SBS-384.

When considering doublet base substitutions or small indels in transcribed regions, for certain mutational events, it is not possible to unambiguously orient these mutations. More specifically, mutations containing both pyrimidine and purine bases cannot be unequivocally attributed to a strand. For example, a TA>AT doublet substitution or a 5’-CATG-3’ deletion cannot be oriented based on the pyrimidine context as both strands contain purine and pyrimidine bases. In contrast, a GG>TT doublet substitution or a 5’-CTTCC-3’ deletion can be oriented as one of the strands is a pure stretch of pyrimidines. Somatic mutations with ambiguous strand orientation have been classified in a separate unknown category (e.g., a TA>AT doublet substitution in a transcribed region is abbreviated as **Q:TA>AT**). In contrast, the classification of somatic indels and DBSs with clear strand orientation has been conducted in a manner similar to the one outlined for single base substitutions.

### Generation of Mutational Matrices and Additional Features

Prior to performing analyses, the tool requires installing a reference genome. By default, the tool supports five reference genomes and allows manually installing any additional reference genome. Installing a reference genome removes the dependency for connecting to an external database, allows for quick and simultaneous queries to retrieve information for sequence context and transcriptional strand bias, and increases the overall performance of the tool.

After successful installation, SigProfilerMatrixGenerator can be applied to a set of files containing somatic mutations from different samples. The tool supports multiple commonly used input formats and, by default, transforms the mutational catalogues of these samples to the above-described mutational matrices and outputs them as text files in a pre-specified output folder.

In addition to generating and plotting matrices from mutational catalogues, SigProfilerMatrixGenerator allows examining patterns of somatic mutations only in selected regions of the genome. The tool can be used to generate mutational matrices separately for: each individual chromosome, for the exome part of the genome, and for custom regions of the genome specified by a BED file. SigProfilerMatrixGenerator can also perform statistical analysis for significance of transcriptional strand bias for each of the examined samples with the appropriate corrections for multiple hypothesis testing using the false discovery rate (FDR) method. Overall, the tool supports the examination of significantly more mutational matrices than prior tools (Table 1) while still exhibiting a better performance (Fig. 3).

**Figure 3:**
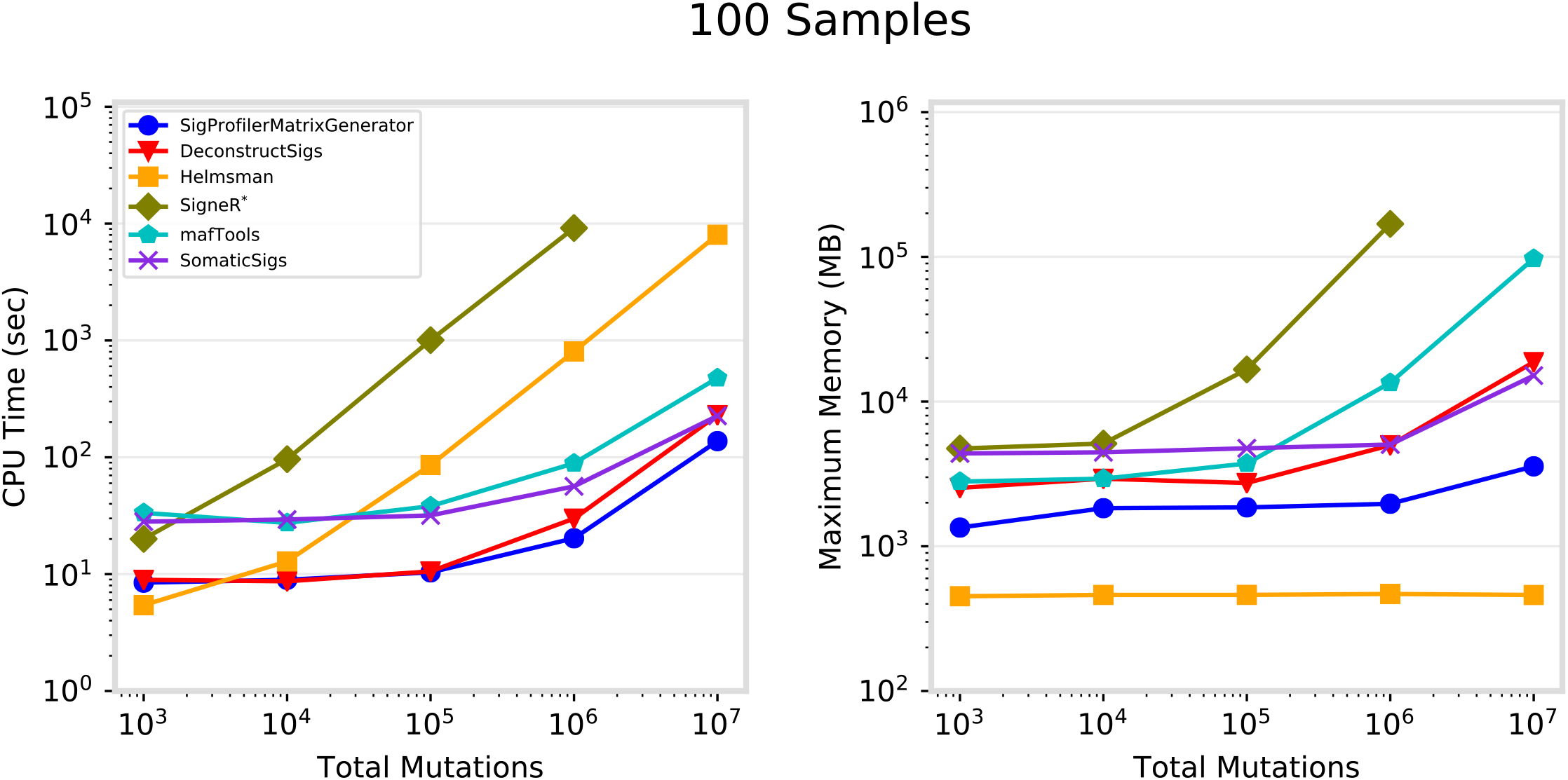
Performance for matrix generation across six commonly used tools. Each tool was evaluated separately using 100 VCF files, each corresponding to an individual cancer genome, containing total somatic mutations between 1000 and 10 million. ***A)*** CPU runtime recorded in seconds (log-scale) and ***B)*** maximum memory usage in megabytes (log-scale). *SigneR was unable to generate a matrix for 10^7^ mutations as it exceeded the available memory of 192GB. Performance metrics exclude visualization.

### Computational Optimization

In addition to its extensive functionality (Table 1), the performance of SigProfilerMatrixGenerator has been optimized for analysis of large mutational datasets. More specifically, as part of the installation process, each chromosome of a given reference genome is pre-processed in a binary format to decrease subsequent query times. This pre-processing reduces a genomic base-pair to a single byte with binary flags that allow immediately identifying the reference base, its immediate sequence context, and its transcriptional strand bias. A single binary file is saved for each reference chromosome on the hard-drive; note that these binary files have similar sizes to ones of FASTA files containing the letter sequences of chromosomes.

When SigProfilerMatrixGenerator is applied to a set of input files, the tool first reformats all input files into a single file per chromosome sorted by the chromosomal positions, e.g., for a human reference genome a total of 25 files are generated: 22 files are generated for the autosomes, two files for the sex chromosomes, and one file for the genome of the mitochondria. Then, the tool processes the input data one chromosome at a time. For example, for a human reference genome, it first loads the reference binary file for chromosome one (~250 megabytes) and all mutations located on chromosome one across all samples are assigned to their appropriate bins in the most extensive classification (e.g., SBS-6144 for single base substitutions). Note that the binary pre-processing of the reference chromosomes makes this a linear operation with identifying the appropriate category for each mutation being a simple binary check against a binary array. After processing all mutations for a particular chromosome, the tool unloads the chromosomal data from memory and proceeds to the next chromosome. When all chromosomes have been processed, the most extensive classification is saved and iteratively collapsed to all other classifications of interests. For example, for single base substitutions, the SBS-6144 is first saved on the hard-drive and then collapsed to SBS-1536 and SBS-384. Then, SBS-1536 and SBS384 are saved on the hard-drive and collapsed, respectively, to SBS-96 and SBS-24. Similarly, SBS-96 and SBS-24 are saved on the hard-drive with SBS-24 being also collapsed to SBS-6, which is also recorded on the hard-drive. Overall, the computational improvements in SigProfilerMatrixGenerator rely on binary pre-processing of reference genomes, iterative analysis of individual chromosomes, and iterative collapsing of output matrices. These computational improvements have allowed computationally outperforming five other commonly used tools.

## Results

The performance of SigProfilerMatrixGenerator was benchmarked amongst five commonly used packages: deconstructSigs [27], mafTools [28], SomaticSignatures [29], signeR [30], and Helmsman [31]. While some of these packages can perform various additional tasks (e.g., extraction/decomposition of mutational signatures), the benchmarking considered only the generation of mutational matrices. The performance was evaluated by measuring the CPU time and maximum memory necessary to generate mutational matrices based on randomly generated VCF files for 100 samples (one file per sample) with different total numbers of somatic mutations: 10^3^, 10^4^, 10^5^, 10^6^, and 10^7^. To maintain consistency, each test was independently performed on a dedicated computational node with an Intel® Xeon® Gold 6132 Processor (19.25M Cache, 2.60 GHz) and 192GB of shared DDR4-2666 RAM. In all cases, the tools generated identical SBS-96 matrices.

In addition to generating an SBS-96 matrix, SigProfilerMatrixGenerator also generates another twelve matrices including ones for indels and doublet base substitutions (Table 1). In contrast, all other tools can only generate a single mutational matrix exclusively for single base substitutions (Table 1). While offering additional functionality, SigProfilerMatrixGenerator exhibits an optimal performance and, in almost all cases, outperforms other existing tools (Fig. 3A). For example, for more than one million mutations, the tool is between 1.5 and 2 times faster compared to the next fastest tool, deconstructSigs. With the exception of Helmsman, SigProfilerMatrixGenerator requires less memory than any of the other tools making it scalable to large numbers of somatic mutations (Fig. 3B). Helmsman’s low memory footprint comes at a price of a significantly slower performance for larger datasets (Fig. 3A).

Lastly, we evaluated whether the exhibited performance is independent of the number of samples by comparing the tools using a total of 100,000 somatic mutations distributed across: 10, 100, and 1000 samples (Supp. Fig 1). SigProfilerMatrixGenerator, deconstructSigs, Helmsman, and mafTools demonstrated an independence of sample number with respect to both CPU runtime and maximum memory usage. The memory usage of SomaticSigs is independent of sample count, however, the runtime increases linearly with the number of samples. The runtime of SigneR is somewhat independent of sample count, however, the memory increases linearly with the number of samples.

## Discussion

SigProfilerMatrixGenerator transforms a set of mutational catalogues from cancer genomes into fourteen mutational matrices by utilizing computationally and memory efficient algorithms. Indeed, in almost all cases, the tool is able to outperform other tools that generate only a single mutational matrix. SigProfilerMatrixGenerator also provides an extensive plotting functionality that seamlessly integrates with matrix generation to visualize the majority of output in a single analysis (Fig. 4). In contrast, most other tools have plotting capabilities solely for displaying an SBS-96 matrix (Table 1). Currently, SigProfilerMatrixGenerator supports only classifications of small mutational events (i.e., single base substitutions, doublet base substitutions, and small insertions and deletions) as we have previously demonstrated that these classifications generalize across all types of human cancer [14]. While classifications for large mutational events (e.g., copy-number changes and structural rearrangements) have been explored by us and others [24, 32, 33] such classifications have been restricted to individual cancer types and it is unclear whether they will generalize in a pan-tissue setting.

**Figure 4:**
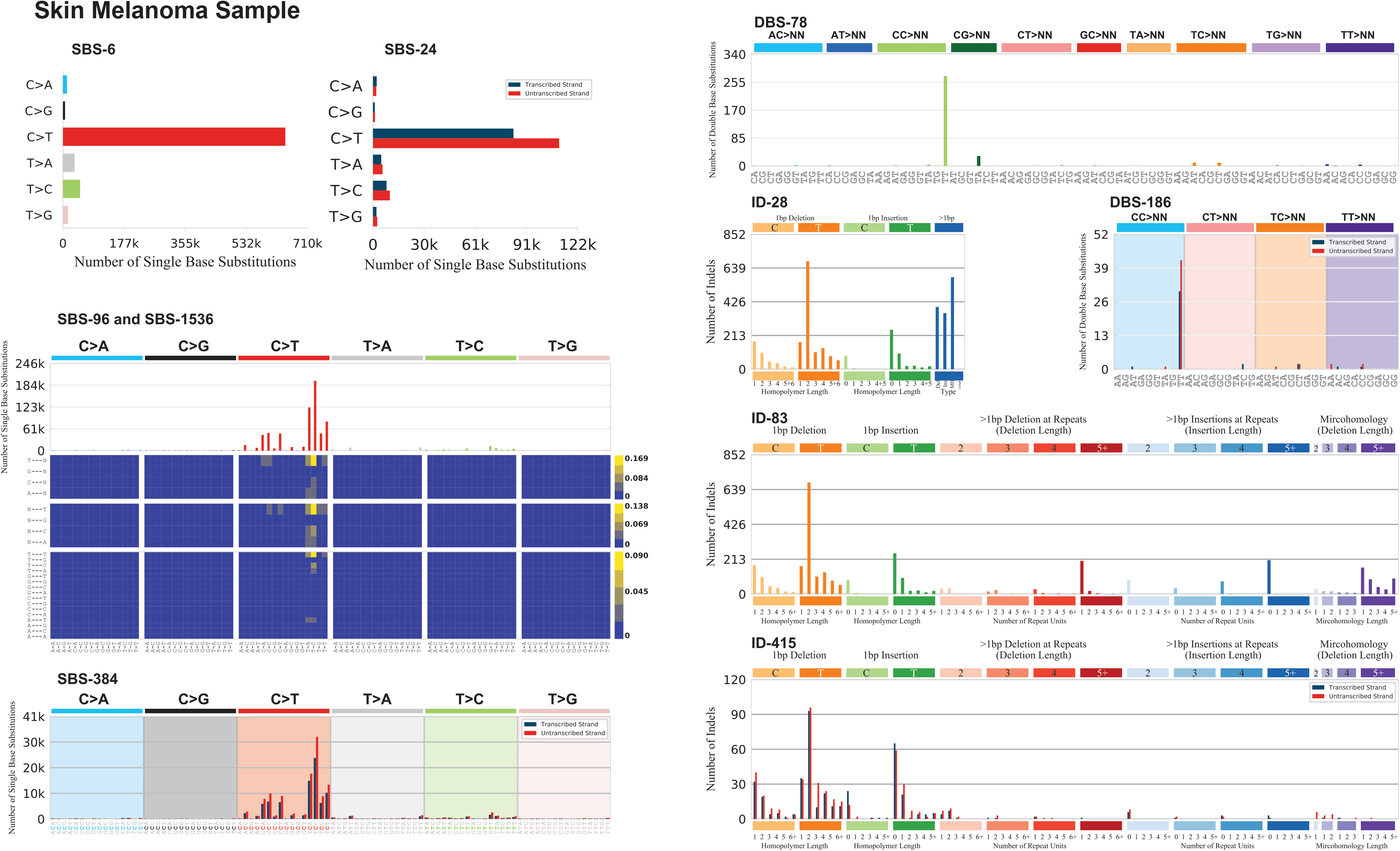
Portrait of a cancer sample. SigProfilerMatrixGenerator provides a seamless integration to visualize the majority of generated matrices. One such functionality allows the user to display all mutational plots for a sample in a single portrait. The portrait includes displaying of each of the following classifications: SBS-6, SBS-24, SBS-96, SBS-384, SBS-1536, DBS-78, DBS-186, ID-28, ID-83, and ID-415. Each of the displayed plots can also be generated in a separate file. Detailed documentation explaining each of the plots can be found at: https://osf.io/2aj6t/wiki/home/.

Importantly, SigProfilerMatrixGenerator is not a tool for analysis of mutational signatures. Rather, SigProfilerMatrixGenerator allows exploration and visualization of mutational patterns as well as generation of mutational matrices that subsequently can be subjected to mutational signatures analysis. While many previously developed tools provide support for examining the SBS-96 classification of single base substitutions, SigProfilerMatrixGenerator is the first tool to provide extended classification of single base substitutions as well as the first tool to provide support for classifying doublet base substitutions and small insertions and deletions.

## Conclusions

A breadth of computational tools was developed and applied to explore mutational patterns and mutational signatures based on the SBS-96 classification of somatic single base substitutions. While the SBS-96 has yielded significant biological insights, we recently demonstrated that further classifications of single base substitutions, doublet base substitutions, and indels provide the means to better elucidate and understand the mutational processes operative in human cancer. SigProfilerMatrixGenerator is the first tool to provide an extensive classification and comprehensive visualization for all types of small mutational events in human cancer. The tool is computationally optimized to scale to large datasets and will serve as foundation to future analysis of both mutational patterns and mutational signatures. SigProfilerMatrixGenerator is freely available at https://github.com/AlexandrovLab/SigProfilerMatrixGenerator with an extensive documentation at https://osf.io/s93d5/wiki/home/.

## Availability and Requirements

### Project name

SigProfilerMatrixGenerator

### Project home page

https://github.com/AlexandrovLab/SigProfilerMatrixGenerator

### Operating system(s)

Unix, Linux, and Windows

### Programming language

Python 3; R wrapper

### Other requirements

None

### License

BSD 2-Clause "Simplified" License

### Any restrictions to use by non-academics

None

## Declarations

### Availability of data and materials

Data sharing is not applicable to this article as no datasets were generated or analyzed during the current study.

### Authors’ contributions

E.N.B. developed the Python and R code and wrote the manuscript. M.N.H., U.M., and M.B. tested and documented the code. M.R.S., S.G.R, and L.B.A. developed the different mutational classifications. L.B.A. supervised the overall development of the code and writing of the manuscript. All authors read the manuscript.

### Ethics approval and consent to participate

Not applicable.

### Consent for publication

Not applicable.

## Acknowledgements

None.

