## Supplementary Figures for "SigProfilerMatrixGenerator: a tool for visualizing and exploring patterns of small mutational events"

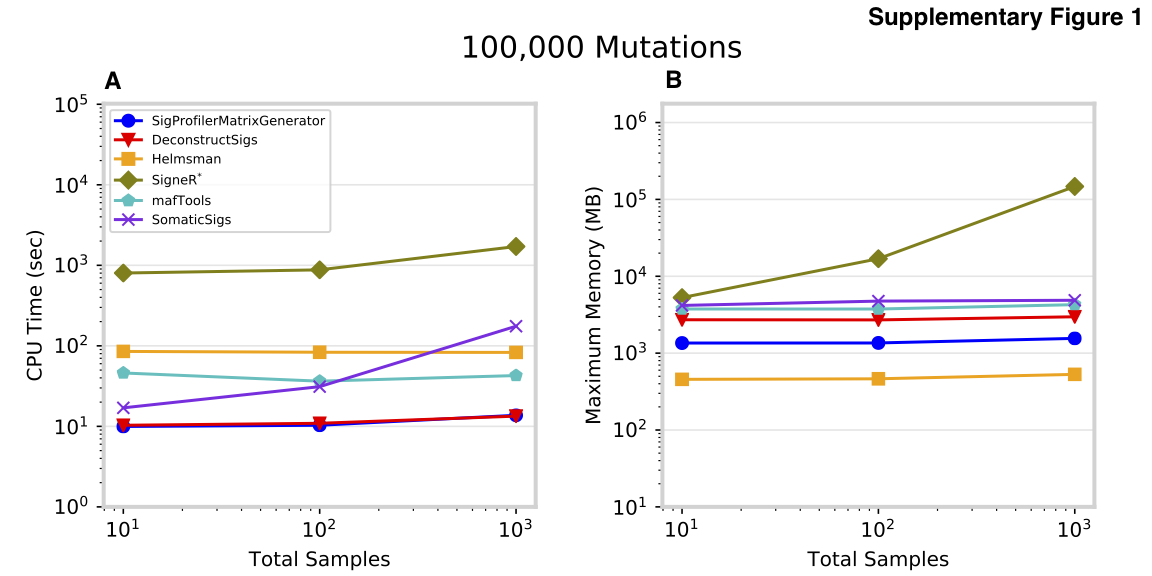

**Supplementary Figure 1: Performance for matrix generation across six commonly used tools.** Each tool was evaluated separately using 10, 100, and 1,000 VCF files, each corresponding to an individual cancer genome, containing a total of 100,000 somatic mutations ***A)*** CPU runtime recorded in seconds (log-scale) and ***B)*** maximum memory usage in megabytes (log-scale). Performance metrics exclude visualization.
